# A DNA Contact Map for the Mouse *Runx1* Gene Identifies Novel Hematopoietic Enhancers

**DOI:** 10.1101/147793

**Authors:** Judith Marsman, Amarni Thomas, Motomi Osato, Justin M. O’Sullivan, Julia A. Horsfield

## Abstract

The transcription factor Runx1 is essential for definitive hematopoiesis, and the *RUNX1* gene is frequently translocated or mutated in leukemia. *Runx1* is transcribed from two promoters, P1 and P2, to give rise to different protein isoforms. Although the expression of Runx1 must be tightly regulated for normal blood development, the mechanisms that regulate *Runx1* isoform expression during hematopoiesis remain poorly understood. Gene regulatory elements located in non-coding DNA are likely to be important for *Runx1* transcription. Here we use circular chromosome conformation capture sequencing to identify DNA interactions with the P1 and P2 promoters of *Runx1*, and the previously identified +24 enhancer, in the mouse multipotent hematopoietic progenitor cell line HPC-7. The active promoter, P1, interacts with nine non-coding regions that are occupied by transcription factors within a 1 Mb topologically associated domain. Eight of nine regions function as blood-specific enhancers in zebrafish. Interestingly, the +24 enhancer interacted with multiple distant regions on chromosome 16, indicating it may regulate the expression of additional genes. The *Runx1* DNA contact map identifies connections with multiple novel hematopoietic enhancers that are likely to be involved in regulating *Runx1* expression in hematopoietic progenitor cells.

## Background

Runx1 is a key regulator of hematopoietic development. Deletion of *Runx1* in mouse embryos is lethal in embryonic stage (E) 12.5 due to the complete absence of definitive blood cell progenitors accompanied by extensive hemorrhaging^1,2^. Runx1 is crucial for hematopoietic stem cell (HSC) emergence and maintenance during development^3^, since conditional ablation of *Runx1* in adult mice results in HSC exhaustion^4^. In acute myeloid leukemia (AML) and myelodysplastic syndrome, RUNX1 function is frequently altered through mutations or translocations^5^, resulting in dysregulation of its target genes. While mutations directly affecting the RUNX1 protein are common in leukemia, mutations in regulatory elements that affect *RUNX1* expression remain enigmatic. As yet unidentified mutations in regulatory elements, such as enhancers, could alter *Runx1* expression, resulting in abnormal hematopoiesis.

*Runx1* is transcribed from two promoters, P1 and P2, to give rise to different protein isoforms^6^. Expression of these isoforms is tightly controlled during hematopoiesis. At the onset of mouse hematopoiesis (E7.5), preceding the generation of HSCs, expression of the P2 isoform(s) is predominant^7,8^. P1 is expressed soon after P2, and its expression is synchronized with the generation of HSCs^7,9^. P1 expression is predominant in the mouse fetal liver, the main site of definitive hematopoietic stem/progenitor cell (HSPC) development from E12.5 onward^9^.

Regulatory elements, such as enhancers, can control the expression of genes via long-range chromatin interactions^10^. One previously identified *Runx1* enhancer is located 24 kb downstream of the P1 transcriptional start site^11,12^. The +24 enhancer (also known as the +23^13^, or the +23.5^12^ enhancer) is active in HSCs that express *Runx1* during mouse embryogenesis^11-13^. The human equivalent to the +24 enhancer (+32 kb downstream of P1) directly contacts the promoters of *RUNX1* in leukemia cell lines^14^. In addition to the +24 enhancer, putative regulatory elements for *RUNX1* have been identified upstream of *RUNX*-P1 and between P1 and P2; however, whether they directly contact the *RUNX1* promoters has not been investigated^15,16^.

Here we used circular chromosome conformation capture sequencing (4C-seq) to identify regulatory elements that interact with an active *Runx1* P1 promoter, versus an inactive P2 promoter. While Hi-C provides genome-wide contact profiles, and Capture Hi-C enriches for interactions with preselected genomic features (usually promoters), 4C-seq can generate very high resolution contact profiles from ‘baits’ of particular interest^17^. Therefore, 4C-seq can yield richer information about a selected genomic region than Hi-C or Capture Hi-C. HPC-7 is a well characterized mouse HSPC line with genomic annotations, including transcription factor (TF) binding, histone modifications and chromatin accessibility^18-20^. 4C-seq in HPC-7 cells identified nine new hematopoietic enhancers that interact with the P1 promoter and +24 enhancer, and that are occupied by hematopoietic TFs. Eight of these were active in zebrafish hematopoiesis. Further, the +24 enhancer was highly interactive both within a topologically associated domain (TAD) harboring *Runx1*, as well as with loci outside the TAD. Collectively, our results point to the formation of a local ‘active chromatin hub’ controlling *Runx1* expression in hematopoietic cells.

## Results

We first confirmed that P1 is actively expressed in HPC-7 cells, while P2 is silent (Supplementary Fig. S1). 4C-seq in HPC-7 cells using the P1 and P2 promoters and the +24 enhancer as ‘baits’ identified genomic interactions at *Runx1* (Fig. 1A). Bait locations were designed taking into account cohesin and CTCF binding sites near both promoters and the +24 enhancer. 4C baits were designed to regions of interest (P1, +24, P2), allowing for comparison of interactions between the active P1 promoter and inactive P2 promoter, with secondary baits located at nearby cohesin/CTCF (cc) binding sites (P1cc, +24cc, P2cc) (Fig. 1A).

**Figure 1.**
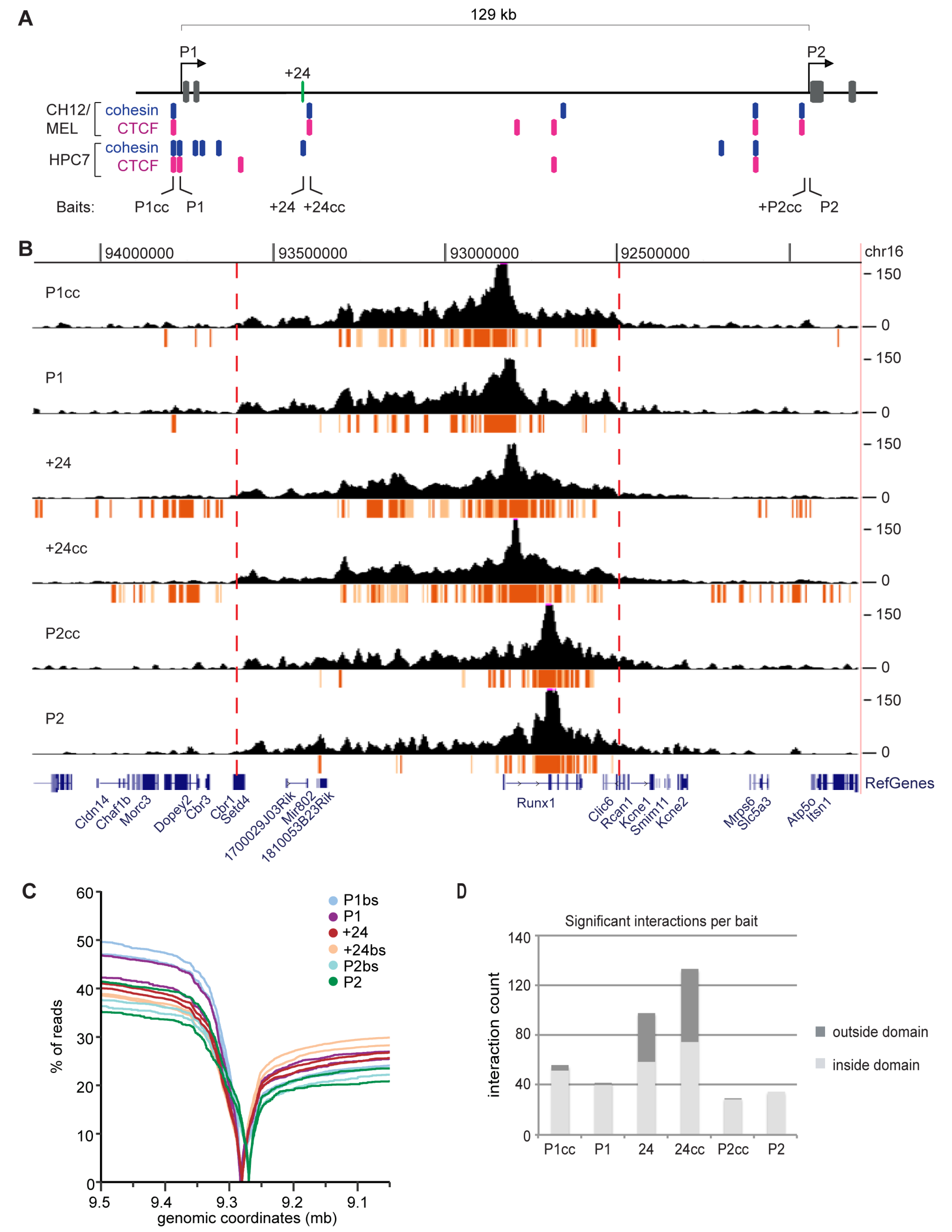
*Runx1* interaction profile reflects the activity of the P1 and P2 promoters and the +24 enhancer. (A) Location of 4C baits at the mouse *Runx1* gene (mm10). Baits are P1cc, P1, +24, +24cc, P2cc and P2; ‘cc’ – cohesin and CTCF binding site. ChIP-seq data of Rad21 (blue) and CTCF (pink) in both MEL and CH12 cells were obtained from ENCODE^50^ and in HPC-7 cells from^18,19^. Exons are in gray and the +24 enhancer in green. (B) 4C-seq profile (read per million normalized running mean of nine successive *Dpn*II digestion fragments) for one replicate per bait with significant interactions (the top fifth percentile of interactions with a FDR of <0.01) that overlap in both replicates shown in a ~2.5 Mb region surrounding the *Runx1* gene. The most highly significant interactions (see Methods) are in red and other significant interactions in orange. Red dotted lines indicate the location of estimated domain boundaries. (C) Read distribution for both replicates of each bait. Line graphs represent the cumulative percentage of reads on chromosome 16 (y-axis) versus the distance from the bait (x-axis), shown in genomic coordinates on chromosome 16 (genome version mm10). The 0 value of each line indicates the location of each bait. (D) The number of significant interactions that overlap in both replicates inside (light grey) and outside (dark grey) the domain for each bait.

### 4C-seq in HPC-7 cells identifies a 1.1 Mb domain harboring *Runx1*

For each bait, two replicate 4C-seq libraries and one control library were sequenced. Reads were predominantly located within a 1.1 Mb region surrounding the *Runx1* gene (Fig. 1B-C), representing a TAD harboring *Runx1*. Domain boundaries are present at the *Cbr1*/*Setd4* genes upstream of *Runx1*, and *Clic6* downstream (Fig. 1B). A comparison with existing Hi-C data from mouse embryonic stem cells, mouse CH12 (erythroleukemia) cells, human GM12878 (lymphoblastoid), human K562 (myeloid leukemia) and human IMR90 (foetal lung) cells revealed that TAD boundaries are conserved, and are consistent with our 4C data (Supplementary Fig. S2). Most P1 and +24 interactions take place upstream of the *Runx1* gene (Fig. 1B,C). In contrast, there are fewer upstream interactions from P2, while downstream interactions are retained (Fig. 1B,C). There are no other coding genes within the *Runx1* domain; however, three inactive non-coding genes^19^, *Mir802* and the long non-coding RNAs *1810053B23Rik* and *1700029J03Rik*, are located near the upstream border of the TAD.

Interactions from ‘cc’ baits (cohesin/CTCF binding sites) were similar to their nearby corresponding baits at P1, +24 and P2 (Fig. 1B). We investigated whether significant interactions for the ‘cc’ baits have more overlap with other cohesin or CTCF binding sites than for the non ‘cc’ baits, but did not find any difference (data not shown). The ‘cc’ baits may be too close to the corresponding P1, P2 and +24 baits to resolve unique interactions, although they are separated by at least one restriction fragment.

### Chromatin interactions anchored by the *Runx1* P1 and P2 promoters and +24 enhancer

Most significant interactions for P1, P2 and the +24 enhancer occur within the 1.1 Mb domain (Fig. 1B). There are ~40-50 significant interactions for P1cc/P1 baits, ~60-75 for +24/+24cc baits, and ~30 for P2/P2cc baits (Fig. 1D). Strikingly, the +24 enhancer in particular forms many significant interactions outside the *Runx1* domain (Fig. 1B,D). Many of these long-range interactions are with other genes or gene promoters on mouse chromosome 16 (Fig. 2). Among the genes contacted by +24 are *Erg*, a hematopoietic TF, and *Tiam1* (T-cell lymphoma invasion and metastasis 1), involved in cell adhesion and cell migration. These distant connections indicate that +24 may also regulate other hematopoietic genes in adjacent domains.

**Figure 2.**
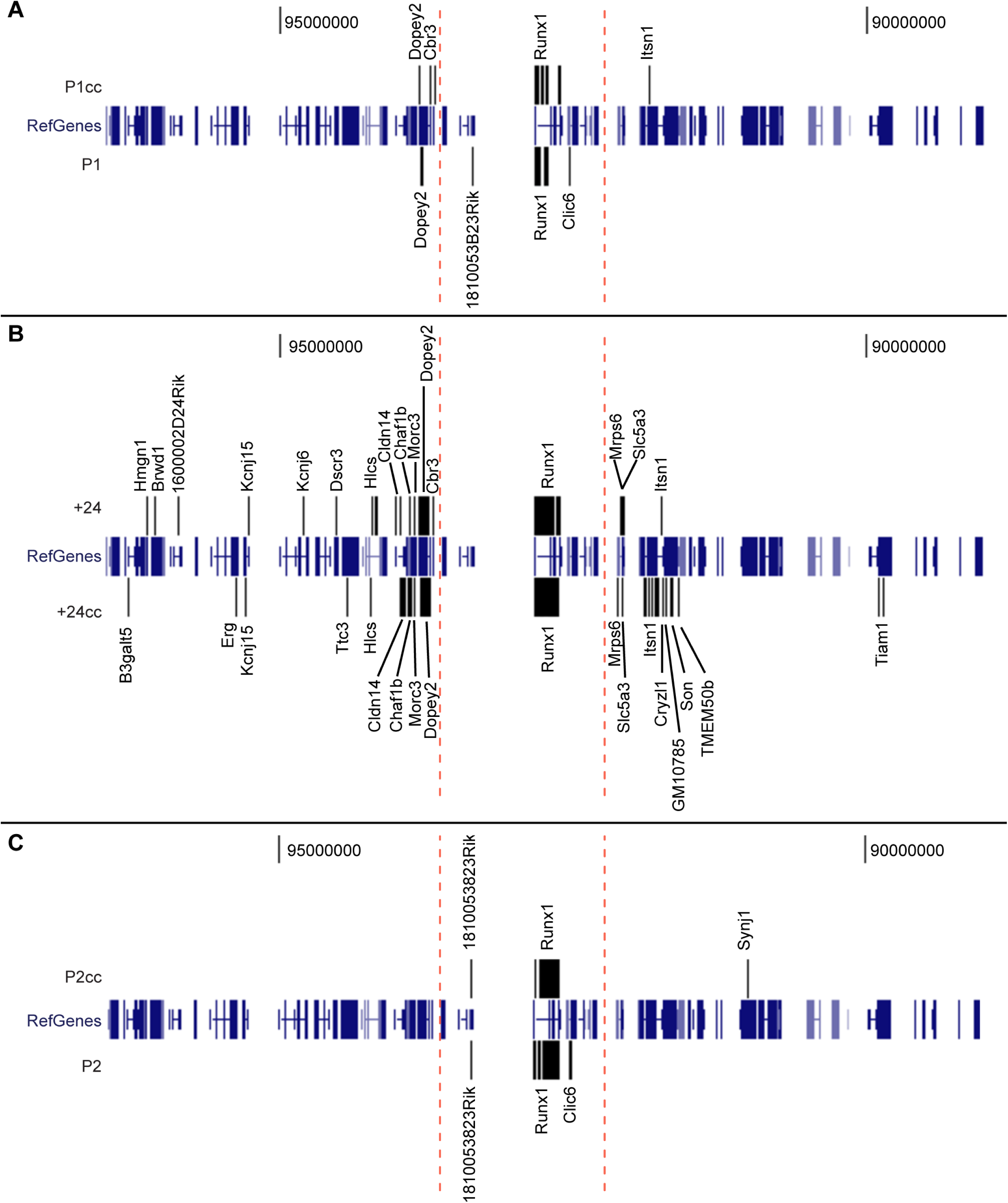
Interactions of the *Runx1* locus with other genes. Significant interactions that overlap with gene bodies or gene promoters (starting from 2 kb upstream of the gene TSS) are shown for the P1cc/P1 (A), +24/+24cc (B) and P2cc/P2 baits (C) (assembly mm10).

### Identification of hematopoietic enhancers

We hypothesized that DNA connections formed from the active P1 promoter and the +24 enhancer may correspond to hematopoietic enhancers that regulate *Runx1* expression. To identify putative enhancers, we aligned significantly interacting sites with the occupancy of thirteen TFs involved in hematopoietic progenitor cell production; enhancer histone modifications and DNase I hypersensitivity sites^18-20^; and conserved non-coding elements (CNE)^11^. We note that the +24 enhancer binds all thirteen hematopoietic progenitor TFs in HPC-7 cells (Fig. 3).

**Figure 3.**
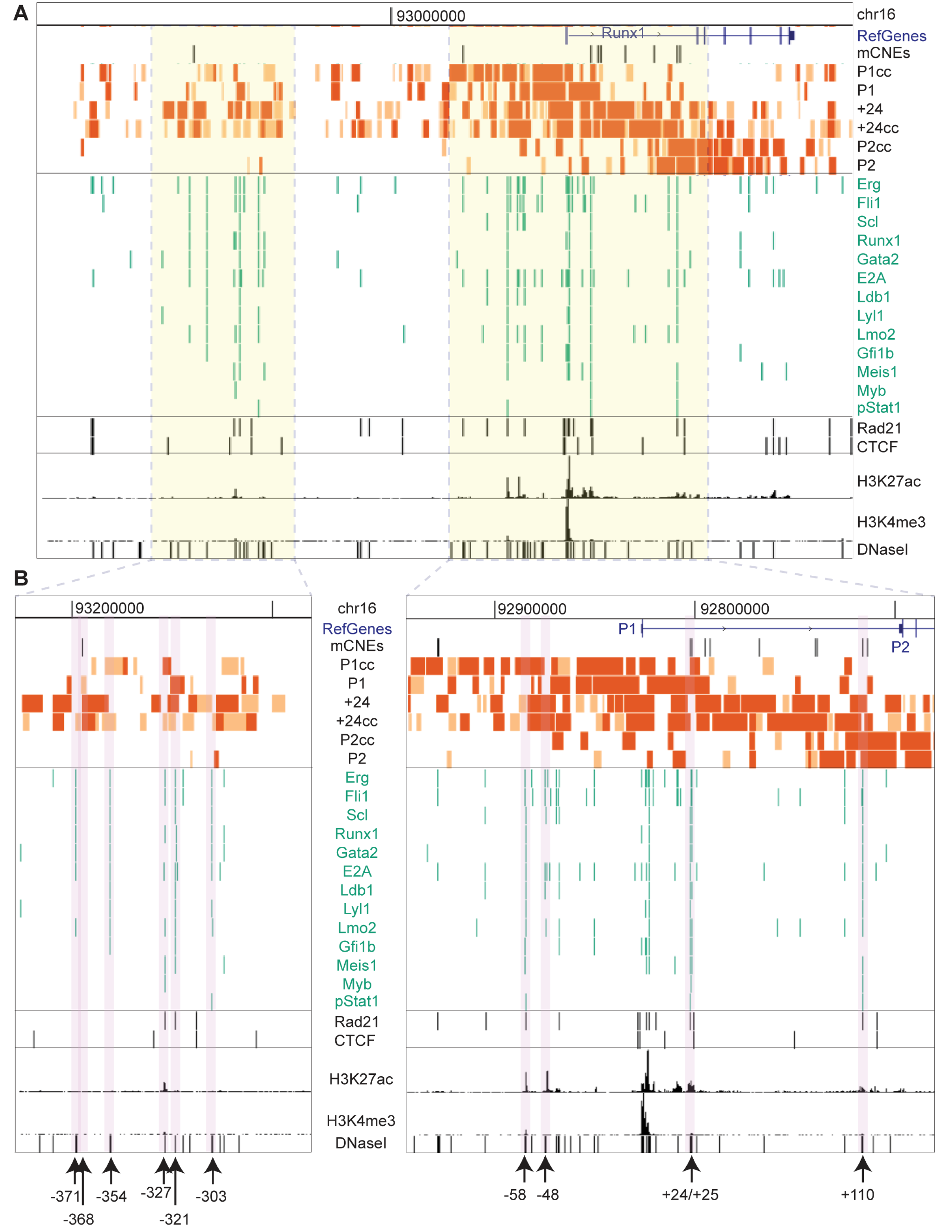
The active P1 promoter and +24 enhancer interact with clusters of hematopoietic transcription factor binding sites. Significant interactions within the domain for each bait are shown together with reference genes (mm9), CNEs^11^, TF binding sites (green), Rad21 and CTCF binding sites (black), H3K27ac, H3K4me3 and DNaseI hypersensitivity sites in HPC-7 cells^18-20^. A 3 Mb region (A) and two zoomed-in views (B) are shown. Locations of TF binding clusters (arrowed) are named according to their distance from the P1 TSS. See also Supplementary Fig. S4.

We found nine other potential enhancers within the *Runx1* domain that form connections to the baits, each occupied by at least six hematopoietic TFs. These enhancers were named according to their distance from the P1 transcriptional start site: -371, -354, -327, -321, -303, -58, -48, and +110 (Fig. 3B). An additional interacting region at -368, a CNE, was located adjacent to a cluster of hematopoietic TFs (Fig. 3B). The putative enhancers either form distinct blocks upstream of *Runx1*-P1 (from -371 to -303 and -48 to -58), or fall between the P1 and P2 promoters (+24 and +110).

Eight out of nine putative enhancers (-371, -368, -354, -327, -321, -58, -48, and +110) form long-range interactions with the P1 promoter, the +24 enhancer, or both (Fig. 3B and Supplementary Table S1). The exception was -303, which does not interact with P1, but instead with the +24 and the P2 promoter. The +24 enhancer interacts promiscuously within the whole domain. Filtering at maximum stringency (see Methods) showed that the +24 enhancer connects to all putative enhancers and both *Runx1* promoters (Fig. 3B). In contrast, the inactive P2 promoter connects only to -303 and +24 (Fig. 3B and Supplementary Table S1). Due to its proximity to P2, interactions between +110 and P2 cannot be resolved. A model of *Runx1* interactions based on the 4C-seq results can be found in Supplementary Figure S3.

### Long range chromatin interactions at hematopoietic enhancers

Long-range chromatin interactions can be mediated by cohesin and CTCF^21^, and cohesin is involved in transcription regulation at active genes^22^. We found that four of the nine enhancer loci (in addition to the +24 enhancer) coincide with Rad21 (cohesin) binding in the absence of CTCF (Fig. 3B and Supplementary Table S1). This is consistent with the idea that cohesin (but not necessarily CTCF) mediates local DNA-DNA interactions within TADs^21^. All Rad21 binding sites interacted with at least one ‘cc’ bait, therefore cohesin could mediate at least a subset of enhancer-promoter communication events in HPC-7 cells.

We compared the 4C interactions identified with recently published Capture Hi-C data in HPC-7 cells^18^ (Supplementary Fig. S4). Capture Hi-C data was only available for interactions anchored at P1, and has a lower coverage and resolution than our 4C-seq study (an average of ~18,000 reads per promoter for Capture Hi-C with a 6-cutter, and over 1 million reads per bait for 4C-seq with a 4-cutter). The Capture Hi-C study in HPC-7 cells identified 15 P1-interacting regions that were reproduced in our study (Supplementary Fig. S4). All of these are upstream of *Runx1*-P1, and most are within the -371 to -303 enhancer cluster (Supplementary Fig. S4). Therefore, our study provides additional *Runx1*-anchored interactions that were not previously described as connected to *Runx1* promoters (enhancers -58, -48 and +110).

### *In vivo* characterization of hematopoietic enhancers

Enhancer regions interacting with *Runx1* recruit hematopoietic TFs in HPC-7 cells, therefore we determined if these regions act as enhancers *in vivo*. Each of the putative enhancers was tested for the ability to drive tissue-specific GFP expression in zebrafish embryos. Eight out of nine drove GFP expression specifically in the intermediate cell mass and posterior blood island, which are sites of hematopoietic progenitor cell production, at 20-24 hours post-fertilization (hpf) (Fig. 4A,B). The -303 enhancer also expressed GFP in keratinocytes, particularly after 24 hpf (Fig. 4C).

**Figure 4.**
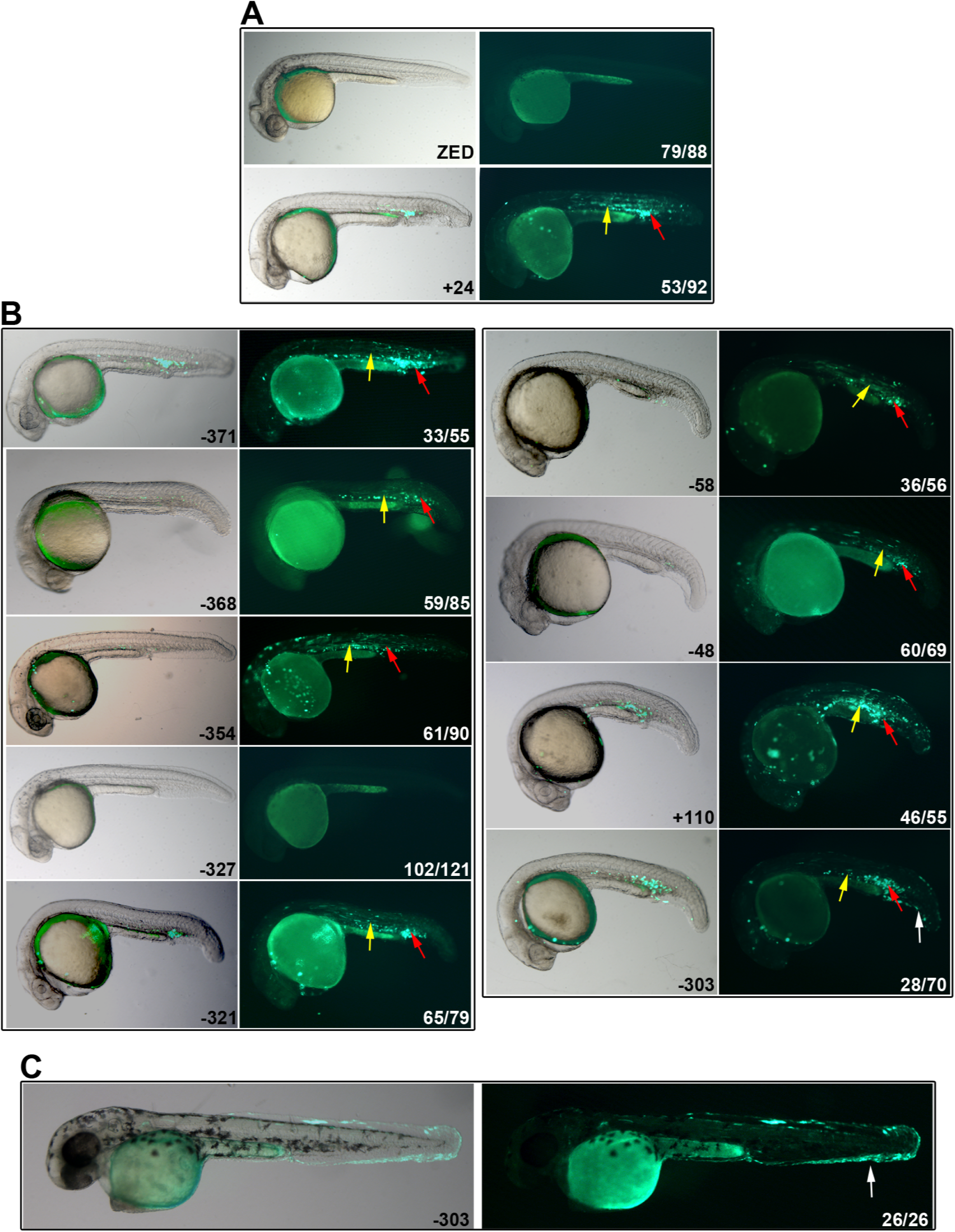
Putative mouse enhancers are active in hematopoietic regions in zebrafish. Whole mount representative lateral images of zebrafish embryos (20-24 hpf [A, B] and ~48 hpf [C]) that were injected with enhancer-GFP plasmids at the one-cell stage; left hand panels are merges with bright field, right hand panels are GFP fluorescence. (A) negative control ZED plasmid shows zero (image) or non-specific fluorescence, while positive control (+24) shows GFP expression in hematopoietic cells of the posterior blood island (red arrows) and intermediate cell mass (yellow arrows). (B) enhancer activity of +110, -48, -58, -303, -321, -327, -354, -368, -371. (C) The -303 enhancer also expressed GFP in keratinocytes (white arrow). The numbers in the right hand panels represent the number exhibiting the representative phenotype out of the total number of fluorescent embryos analyzed.

Interestingly, -303 was the only enhancer identified that interacted with the P2 promoter of *Runx1*, rather than P1. Despite the occupancy of multiple hematopoietic TFs in HPC-7 cells, and the presence of similar TF binding motifs compared to the other enhancers (Fig. 3 and Supplementary Table S1-2), we did not observe enhancer activity for -327.

In summary, we have assigned *in vivo* function to multiple putative enhancers upstream of *Runx1* that were previously identified *in silico*^18^. These regions not only drive hematopoietic expression, but also physically connect with the *Runx1*-P1 promoter, lending confidence to the concept that they are *bona fide* regulators of *Runx1* transcription.

## Discussion

4C analysis in HPC-7 cells generated a high-resolution connectivity map of the genomic region harboring *Runx1*, and confirmed previously identified upstream connections from P1 to putative regulatory elements^18^. *Runx1* appears to be contained within a ~1 Mb chromatin domain, consistent with Hi-C analyses in other cell types^23,24^. We observed multiple connections both up- and downstream from all 4C-seq baits (*Runx1*-P1, *Runx1*-P2 and +24 enhancer). Significantly, there were many more upstream connections anchored by the active elements (*Runx1*-P1 and +24).

Surprisingly, the +24 enhancer forms many up- and downstream connections outside of the *Runx1* TAD. These comprise up to one-third of all connections formed and include other hematopoietic genes, such as *Tiam1* downstream, and *Erg* upstream. *Erg* and *Tiam1* dysregulation is associated with several tumor types, including AML and B- and T-cell lymphomas^25,26^. Strikingly, these genes are located over 2 Mb away from *Runx1*. These findings raise the interesting possibility that the +24 enhancer acts as a scaffold to recruit multiple promoters, enhancers and TFs over long distances in cis. Although the +24 enhancer is a specific marker of HSPCs, not all HSPCs that are marked by expression from this enhancer also express *Runx1*^27^. Therefore, our results are consistent with the idea that the +24 enhancer regulates genes in addition to *Runx1.* Connections to +24 identified by our 4C-seq may point to the identity of some of these genes.

Our 4C-seq data identified multiple chromatin connections that intersect with some of the strongest indicators of upstream enhancers characterized by the binding of clusters of TFs and epigenetic modifications^18,20^. Eight out of nine of these DNA regions are able to drive hematopoietic expression in zebrafish, strongly suggesting an *in vivo* function. Two enhancers termed -59 and +110 were previously shown to drive *LacZ* expression in mice^15^; these are likely to be equivalent to the -58 and +110 enhancers identified here. Our data show in addition that these regions are in contact with P1 and the +24 enhancer.

The -303 enhancer interacts with P2 rather than P1 and, in addition to being active in hematopoietic sites, it drives expression in keratinocytes, indicating that it could act in a complex that keeps P2 silent; and/or that it regulates expression of *Runx1* in other tissues. Interestingly, *Runx1* is actively expressed in mouse keratinocytes where it is important for hair follicle development^28^. Neuronal cells also express *Runx1*^29^, and there may be additional regulatory elements that control *Runx1* expression in a manner distinct from hematopoietic expression. In support of this idea, we previously determined that cohesin and CTCF influence *runx1* expression in hematopoietic, but not neuronal cells, in zebrafish^30,31^.

Cohesin and CTCF organize chromatin structure and when present in combination, they appear to negatively correlate with the HSPC TFs in HPC-7 cells^18^. However, we observed a coincidence of cohesin subunit Rad21 binding in the absence of CTCF with four out of nine identified enhancers, as well as the +24 enhancer. This suggests that CTCF-independent cohesin mediates a subset of enhancer-promoter looping in combination with TFs. This interpretation is consistent with previously identified CTCF-independent functions for cohesin in genome organization and transcription^32^. Importantly, cohesin mutations are prevalent in AML and other myeloid malignancies^33^, and are categorized together with *RUNX1* and spliceosome mutations in a genetic category that confers poor prognosis in AML^5^. Cohesin mutations led to increased chromatin accessibility of *Runx1* as measured by ATAC-seq^34^, raising the possibility that spatiotemporal regulation of *Runx1* is cohesin-dependent in mouse and human, as was previously observed in zebrafish^30^.

4C-seq in HPC-7 cells has provided new high resolution connectivity data that sheds light on the genomic organization of *Runx1*, an important hematopoietic transcription factor. These data confirm and extend previous analyses, and furthermore, provide insight into the function of enhancers that have potential to regulate *Runx1* expression. The data presented here set the scene for functional analyses to precisely determine how *Runx1* is regulated, including CRISPR/Cas9-mediated interference with enhancer activity. They also provide a rationale for screening patients with myeloproliferative disorders for mutations in enhancer regions.

## Methods

### Cell culture

HPC-7 cells were maintained at a density of 1-10 x 10^5^ cells/mL in Iscove’s modified Dulbecco’s media (Gibco®) supplemented with 3.024 g/L sodium bicarbonate, 10% fetal bovine serum (Moregate, New Zealand), 10% stem cell factor conditioned media and 0.15 mM monothiolglycerol (Sigma-Aldrich) as previously described^35^. SCF-conditioned media was obtained from culturing BHK-MKL cells maintained in Dulbecco’s modified Eagle media (Sigma-Aldrich) supplemented with 3.5 g/L glucose, 3.7 g/L Sodium Bicarbonate and 10% FBS.

### 4C-seq library preparation

4C library preparation was performed as previously described^36^ with modifications. Three libraries were generated, two replicates (passage 8 and passage 10 cells) and one control (a 1:1 mix of both replicates for which the first ligation step was omitted). Cells were cross-linked in 2% formaldehyde, 5% FBS and 1x PBS for 10 minutes at room temperature while rotating. Formaldehyde was quenched with a final concentration of 125 mM glycine for 5 minutes on ice while inverting several times. Cell pellets were washed twice with ice-cold 1x PBS.

Nuclei were harvested by lysing the cell pellets in ice-cold lysis buffer (10 mM Tris pH 8.0, 10 mM NaCl, 0.2% NP-40, protease inhibitors) for 10 minutes on ice. Nuclei were then resuspended in 1.2x *Dpn*II restriction buffer (New England Biolabs) and 0.3% SDS and incubated for 1 hour at 37°C while shaking. Triton X-100 was then added to a final concentration of 1.8% and the reaction was left at 37°C while shaking for another hour. Chromatin was digested with 800 U of *Dpn*II overnight at 37°C while shaking. *Dpn*II was inactivated by adding SDS to a final concentration of 1.3% and incubating at 65°C for 20 minutes. Nuclei were diluted into a volume of 7 mL containing 1.01x T4 DNA ligase buffer (Life Technologies) and Triton-X100 at a final concentration of 1% and incubated at 37°C for 1 hour. Ligations were carried out with 100 U of T4 ligase (Life Technologies) for 4.5 hours at 16°C and 30 minutes at room temperature while shaking. For control libraries, ligase was omitted. Samples were proteinase K treated and reverse-crosslinked overnight at 65°C. Samples were then treated with RNase A at 37°C for 30 minutes. DNA was purified by phenol/chloroform extraction and ethanol precipitated.

A second digestion was performed with 25 U of *Bfa*I (New England Biolabs) for P1 and P2 baits or *Mse*I (New England Biolabs) for +24 baits in restriction buffer overnight at 37°C while shaking.

Restriction was inactivated by adding SDS to a final concentration of 1.3% and incubating at 65°C for 20 minutes. A second ligation was performed in the same way as the first ligation, except that ligations were incubated overnight. DNA was purified with two phenol/chloroform extractions and one chloroform extraction followed by ethanol precipitation. DNA concentrations were measured using a Qubit® 3.0 Fluorometer (Life Technologies) and Qubit® double-stranded DNA (dsDNA) High Sensitivity Assay kit (Life Technologies).

For each bait, a total of 1 μg of DNA was amplified by PCR using Q5® High-Fidelity DNA Polymerase (New England Biolabs). Bait primer sequences are listed in Supplementary Table S3. PCR products were purified using the QIAquick PCR Purification kit (QIAGEN). DNA concentrations were measured using a Qubit® 3.0 Fluorometer and average fragment size by 2100 Bioanalyser (Agilent Technologies) using a High Sensitivity DNA Kit (Agilent Technologies). Amplicons from the six different baits were mixed equally based on the concentration, average fragments size and ratio of demultiplexed 4C baits obtained from an initial MiSeq run. Libraries were prepared with Prep2Seq^TM^ DNA Library Prep Kit from Illumina^TM^ (Affymetrix) and TruSeq® adaptors (Illumina). Libraries were mixed equimolarly and sequenced as 125 bp paired-end reads on two Illumina^TM^ HiSeq 2500 lanes by New Zealand Genomics Limited.

### 4C-seq data analysis

4C-seq data was analysed in command-line and the R statistical environment^37^ and visualized using the University of California, Santa Cruz (UCSC) genome browser (http://genome.ucsc.edu/) with mouse assemblies mm9 and mm10^38,39^ and using the R package ggplot2^40^. Baits were demultiplexed based on bait primer sequences up to and including the digestion site using a custom awk script, allowing 0 mismatches. Only read pairs that had the forward and reverse bait sequences in the correct orientation were selected. Adapter sequences, bait sequences up to but excluding the digestion site, and bases with a Phred quality score under 20 were then trimmed from the reads, using the fastq-multx, fastq-mcf and cleanadaptors v1.24 tools^41,42^. Quality of reads was assessed using FastQC (http://www.bioinformatics.babraham.ac.uk/projects/fastqc/)^43^.

Reads with a minimum length of 30 bp were mapped to the mm10 reference genome using Bowtie1^44^, allowing 0 mismatches. Mapped reads were assigned to *Dpn*II digestion fragments using fourSig^45^. The following reads were removed from the files: 1) self-ligated reads, 2) uncut reads (fragments adjacent to baits), and 3) reads at fragments that have at least 1 read in the control (non-ligated) library. The running mean was calculated from the sum of read counts from nine successive fragments, which was obtained using fourSig^45^, and was read per million normalized.

Significant interaction calling was performed using the R package fourSig with the following settings: window size of 3, 1000 iterations, fdr of 0.01, fdr.prob of 0.05 (which selects the top fifth percentile of interactions with a FDR of <0.01), and only included mappable fragments^45^. Significant interactions were called for two regions: 1) the whole of chr16, and 2) from chr16:92,250,000-93,635,000 (within the domain). Significant interactions in both replicates were overlapped using the bedIntersect tool from UCSC^46^.

In the fourSig package, significant interactions can be categorized into three categories: 1) interactions that are significant after the reads from the fragment with the highest read count is removed, 2) interactions that are significant when the fragment with the highest read count is averaged to the read counts of the neighbouring fragments, and 3) interactions that are significant only when all fragment read counts are included^45^. For this study, only category 1 and 2 interactions that overlap between both replicates were included, as they are more likely to represent true interactions (because they span multiple fragments), and were shown to be more reproducible between replicates than single-fragment interactions^45^. Furthermore, we distinguished category 1 interactions that overlap between both replicates from other category 1 and 2 interactions by coloring them red and orange, respectively, to visualise the most significant interactions. For conversion from assembly mm10 to mm9, the liftOver tool from UCSC was used (http://genome.ucsc.edu/)^46^. Gene annotations used in Figures are UCSC reference genes.

### Zebrafish enhancer assay

*Runx1* regulatory regions were amplified from HPC-7 gDNA or from I-SceI-zhsp70 plasmid containing the -368, +24 and +110 sequences^11^ (primer sequences are in Supplementary Table S3) and cloned into the zebrafish enhancer detection vector^47^. Primers amplified the TF binding peak +/-200 bp, except for -321. The -368, +24 and +110 fragments are 471, 529 and 579 bp, respectively^11^. 30 pg vector DNA and 120 pg Tol2 transposase mRNA^48^ was injected into 1-cell zebrafish embryos. Embryos were imaged at 20-24 or ~48 hpf using a Leica M205FA stereomicroscope with a DFC490 camera and LAS software (Leica Microsystems), images were processed using Adobe Photoshop. Zebrafish were maintained as described previously^49^ and zebrafish handling and procedures were carried out in accordance with the Otago Zebrafish Facility

Standard Operating Procedures. The University of Otago Animal Ethics Committee approved all zebrafish research under approval AEC 48/11.

### Identification of conserved non-coding elements

Mouse conserved non-coding elements (mCNEs) were identified as described previously^11^.

### ChIP-seq, Capture Hi-C and DNase I hypersensitivity data

Occupancy of the transcription factors Erg, Fli1, Scl, Runx1, Gata2, E2A, Ldb1, Lyl1, Lmo2, Gfi1b, Meis1, Myb, phospho-Stat1, Pu.1, Stat3, Eto2, Cebp-α, Cebp-β, Elf1, Nfe2, p53, cMyc, Egr1, E2f4, cFos, Mac and Jun; Rad21 and CTCF; H3K27ac and H3K4me3; DNase I hypersensitivity sites; and Capture Hi-C data in HPC-7 cells was obtained from previously published data^18-20^. Rad21, Smc3 and CTCF chromatin immunoprecipitation sequencing (ChIP-seq) data in MEL and CH12 cells were obtained from ENCODE^50^.

## Acknowledgements

HPC-7 and BHK/MKL cells were kindly provided by Kathy Knezevic and John Pimanda. TruSeq® adaptors were kindly provided by Ian Morison. The authors would like to thank Anita Dunbier and Sofie Van Huffel for help with development of 4C protocols, and Noel Jhinku for expert management of the Otago Zebrafish Facility. This research was funded by Royal Society of NZ Marsden Fund [grant number 11-UOO-027 to JAH] and the Health Research Council of NZ [grant number 15/229 to JAH].

## Authorship contributions

JM, AT, MO, JMO, JAH, designed experiments; JM, AT, performed experiments; JM, AT, MO, JMO, JAH analyzed data; JM and JAH wrote the paper with input from the other authors.

## Additional information

### Availability of data

The 4C-seq dataset is accessible through GEO Series accession number GSE86994 (https://www.ncbi.nlm.nih.gov/geo/query/acc.cgi?acc=GSE86994).

